# Shotgun sequencing data and SSR mining data of aibika (*Abelmoschus manihot,* Malvaceae)

**DOI:** 10.1101/2025.11.23.689977

**Authors:** Ronan Rivallan, Andrea Garavito, Floriane Lawac, Nadia Robert, Janet Paofa, Jean-Pierre Labouisse

**Affiliations:** CIRAD, UMR AGAP Institut, F-34398 Montpellier, France; UMR AGAP Institut, Univ Montpellier, CIRAD, INRAE, Institut Agro, Montpellier, France; VARTC, PO Box 231, Santo, Vanuatu; IAC Institut Agronomique Néo-Calédonien, BP 73, 98890 Païta, New Caledonia, France; NARI, Southern Regional Centre, Laloki, PO Box 1828, Port Moresby, Papua New Guinea

**Author notes:** **Corresponding author’s email address**.

**Keywords:** Bele, Genetic diversity, Illumina MiSeq, Island cabbage, Microsatellites, New Caledonia, Papua New Guinea, Vanuatu

## Abstract

Aibika (*Abelmoschus manihot*) is a tropical leafy vegetable with great potential in the prevention of malnutrition. Its high fibre and micronutrient content makes it a valuable complement to staple foods such as rice and starchy crops consumed in South Asia and the Pacific. The present study is the first to report the development of a set of 21 nuclear single sequence repeat (SSR) markers of *A. manihot* from genomic sequence data obtained using NGS technology. The DNA library was prepared from a pool of four aibika accessions and sequenced using an Illumina MiSeq system. A total of 1,295,217 pair-end reads were generated. Raw data are available in the European Nucleotide Archive (accession number PRJEB88210; https://www.ebi.ac.uk/ena/browser/view/PRJEB88210). The sequences were assembled using the ABySS software. In total 651,320 contigs were generated. Using MISA Perl script and Primer3 software, we identified 8014 SSR motifs, of which 4637 had a suitable primer design. The contigs were blasted with NCBI-BLAST on okra reference genome. After selection based on motif type, repeat length, amplicon size and occurrence blast, we retained 91 candidate SSR loci, which were tested on 23 aibika accessions. Finally, we validated 21 high quality SSR loci by genotyping 45 accessions from three Pacific countries. The number of alleles per locus ranged from 3 to 21 with an average of 7.81 alleles/locus. The 21 selected SSRs were found to be useful in discriminating between accessions and revealing the diversity structure of *A. manihot*. They will help to optimize genebanks management and breeding programmes, and guide future collection activities.

## BACKGROUND

*Abelmoschus manihot* (L.) Medik., commonly known as aibika, bele, or island cabbage, is a tropical leafy vegetable that has the potential to prevent malnutrition and micronutrient deficiencies [1–3]. The species is native to the Indomalaya biogeographical realm and has spread to New Guinea, Northern Australia, and Pacific Island countries. It is a popular crop in Melanesia where it is propagated mainly by stem cuttings, although seed propagation has been reported [4]. Research efforts to assess genetic diversity have focused primarily on its congener, *A. esculentus* (L.) Moench (okra) [5–7]. In contrast, our understanding of the genetic diversity of *A. manihot* is limited to a study conducted by Rubian-Yalambing et al. [8] who analyzed 23 aibika accessions of Papua New Guinea with RAPD and DAMD markers. The objective of our study was to develop a set of nuclear simple sequence repeat (SSR) markers for assessing the genetic diversity of *A. manihot*. To achieve this, we sequenced a DNA library with an Illumina MiSeq system. Raw sequencing data were processed using computational pipelines to find candidate SSR loci. Finally, we validated a set of 21 polymorphic SSR markers by genotyping 45 accessions from three Pacific Island countries.

## DATA DESCRIPTION

### Sequencing and genome assembly data

A pooled DNA extract of four *A. manihot* accessions from Vanuatu (VUT084, VUT290, VUT301, VUT313) was used to prepare the DNA library which was sequenced on an Illumina MiSeq system. A total of 1,295,217 pair-end reads were generated. The raw sequences have been deposited in the European Nucleotide Archive (ENA) at the European Bioinformatics Institute (EMBL-EBI) under accession number PRJEB88210.

The sequences were assembled using the ABySS software with a k-mer length of 64. In total 651,320 contigs were generated. The basic statistics of genome assembly are summarized in Table 1.

**Table 1.** Genome assembly statistics of *Abelmoschus manihot*.

| Parameter | Value |
| --- | --- |
| Number of contigs | 651,320 |
| Number of contigs ≥ 500 bp | 812 |
| Sum of contig lengths | 819,639 bp |
| Largest contig | 11,070 bp |
| Shortest contig | 500 bp |
| N25 | 2909 |
| N50 | 1110 |
| N75 | 596 |
| L50 | 157 |

### SSRs data

Using MISA Perl script and Primer3 software, we identified 8014 microsatellite motifs, of which 4637 had a suitable primer design. We excluded complex motifs (4497 retained), then selected amplicon sizes ranging between 100 and 350 bp (384 loci retained). We blasted the selected primer pairs on the okra genome and we kept the primer pairs with only one hit (145 retained). After selection on motif design (from 2 to 6), 96 candidate primer pairs were tested for amplification by genotyping a subset of 23 *A. manihot* samples. Primer pairs were classified according to their polymorphism and the overall quality of the profile. Lastly, we selected 21 polymorphic markers (19 single-locus markers and one two-locus marker) with no ambiguity in allele size determination (Annex 1).

### Genotyping data

Forty-five accessions from three Pacific countries were genotyped using the 21 selected SSR markers. The number of alleles observed per locus and per country is presented in Table 2. For the 21 loci, we found a total of 164 alleles and an average of 7.81 alleles per locus.

**Table 2.** Number of alleles, per locus and per country, observed by genotyping the 45 accessions of *Abelmoschus manihot*.

| Locus | All countries<br>(N=45) | Per country (n = 15) |  |  |
| --- | --- | --- | --- | --- |
|  |  | PNG | NCL | VUT |
| AmCir0203 | 7 | 4 | 5 | 4 |
| AmCir0518 | 6 | 4 | 5 | 2 |
| AmCir0731 | 14 | 10 | 3 | 6 |
| AmCir0868 | 5 | 5 | 4 | 3 |
| AmCir1791 | 3 | 3 | 2 | 2 |
| AmCir1965 | 4 | 3 | 2 | 3 |
| AmCir2957 | 9 | 8 | 3 | 4 |
| AmCir3417 | 7 | 5 | 5 | 5 |
| AmCir3540 | 7 | 4 | 6 | 4 |
| AmCir4539 | 6 | 4 | 4 | 4 |
| AmCir4803 | 21 | 13 | 8 | 6 |
| AmCir5172 | 6 | 6 | 3 | 3 |
| AmCir5434 | 7 | 6 | 4 | 3 |
| AmCir5534 | 8 | 5 | 4 | 4 |
| AmCir5677 | 10 | 8 | 4 | 3 |
| AmCir6454 | 9 | 7 | 4 | 5 |
| AmCir8412 | 9 | 6 | 6 | 4 |
| AmCir8690 | 6 | 4 | 3 | 3 |
| AmCir9461 | 3 | 2 | 2 | 3 |
| AmCir9548a | 13 | 11 | 6 | 7 |
| AmCir9548b | 4 | 3 | 3 | 2 |
| Total (A) | 164 | 121 | 86 | 80 |
| (A)/21 loci | 7.81 | 5.76 | 4.10 | 3.81 |

PNG=Papua New Guinea; NCL=New Caledonia; VUT= Vanuatu

The DARwin software was used to compute the dissimilarity matrix and to perform a factorial analysis. Using the set of 21 markers, we were able to distinguish the *A. manihot* accessions within each genebank and to show how genetic diversity is structured among three countries of the SW Pacific (Fig. 1).

**Fig. 1.**
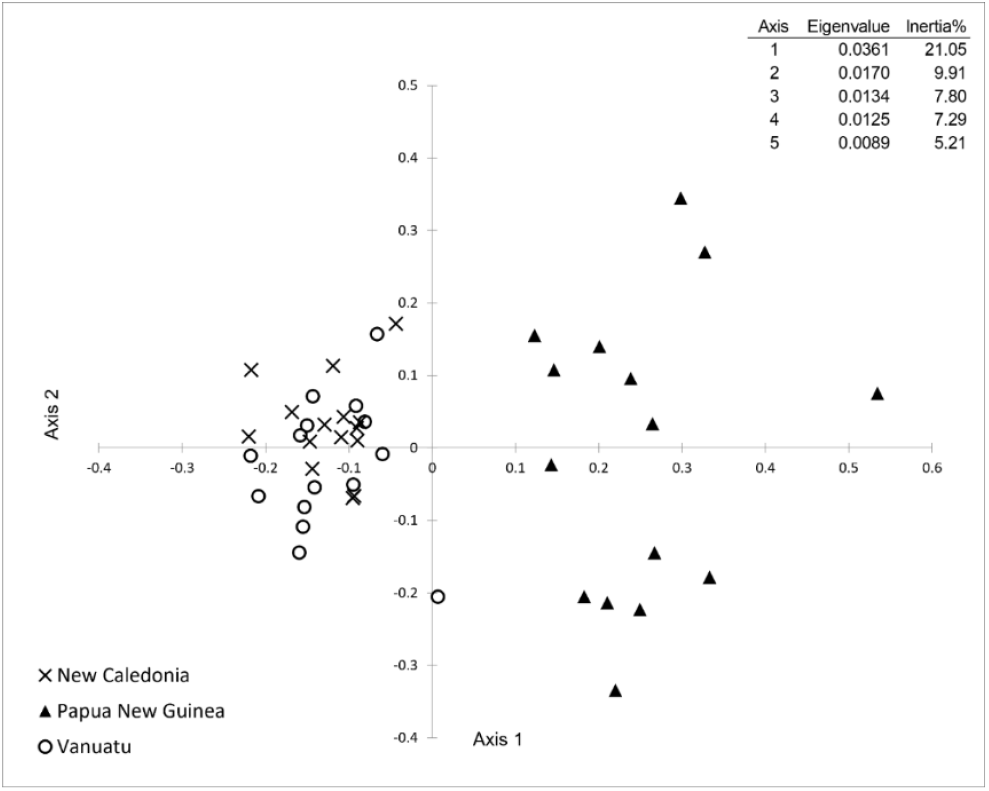
Graphical representation of the factorial analysis (PCoA) of the distance matrix computed from the genotyping data of the 45 accessions of *Abelmoschus manihot*, using the 21 selected markers.

## EXPERIMENTAL DESIGN, MATERIALS AND METHODS

### Plant material sampling

The study screened 45 accessions, maintained by research institutions in three Pacific countries (Papua New Guinea, New Caledonia and Vanuatu), sourced from their respective field genebanks (Table 3). Among them, we selected four accessions from different islands of Vanuatu, exhibiting distinct morphological traits, to prepare the DNA library for sequencing. All of the 45 accessions were genotyped to validate the candidate SSR markers.

**Table 3.** List of the *Abelmoschus manihot* accessions used in the study.

| Provider | Country of origin | Accession numbers |
| --- | --- | --- |
| Institut Agronomique néo-Calédonien (IAC)<br>Port Laguerre station, Païta,<br>New Caledonia | New Caledonia | NCL014; NCL024; NCL037; NCL045;<br>NCL087; NC089; NCL100; NCL113;<br>NCL132; NCL134; NCL136; NCL138;<br>NCL144; NCL146; NCL151 |
| National Agricultural Research<br>Institute (NARI)<br>Bubia and Laloki stations,<br>Papua New Guinea | Papua New Guinea | BUBAm002; BUBAm003; BuBAm006;<br>BUBAm009; BUBAm012; BUBAm013;<br>LALAm039; LALAm200; LALAm206;<br>LALAm223; LALAm239; LALAm253;<br>LALAm254; LALAm257; LALAm260 |
| Vanuatu Agricultural Research<br>and Technical Center (VARTC),<br>Santo, Vanuatu | Vanuatu | VUT006; VUT015; VUT062; VUT063;<br>VUT084*; VUT262; VUT271; VUT273;<br>VUT277; VUT290*; VUT301*; VUT313*;<br>VUT315; VUT349; VUT371 |

\*Accessions used for sequencing

### DNA extraction

We selected one young, healthy green leaf from each sampled plant. A piece of about 20 cm^2^ of young, healthy, green leaf was taken from each sampled plant About 20 cm^2^ of leaf tissue was cut out with scissors and dried in a ventilated oven at 40 °C for 48 hours. The dried samples were sent to the genotyping platform at the Centre de Coopération Internationale en Recherche pour le Développement (CIRAD), Montpellier, France for DNA extraction, sequencing, and genotyping. For each sample, 50 mg of dry leaves were ground 2 times at 1400 rpm during 1 min 30 sec with 2 steel beads of 3 mm on Genogrinder 2010 (Cole-Palmer®, Antylia Scientific, Vernon Hills, IL, USA), incubated at 56 °C for 45 min with 600 µL Lysis Buffer CF (Macherey-Nagel, #740946, Düren, Germany) and 200 ng Proteinase K (Macherey-Nagel, #740506). DNA was then purified on Thermo Scientific KingFisherTM Duo Prime System (Thermo Fisher Scientific, Watham, MA, USA) with NucleoMag® C-Beads (Macherey-Nagel, #744401). The quality of the genomic DNA was checked using 1% agarose gel (1 g/100 mL) electrophoresis and a Nanodrop 2000c spectrophotometer (Thermo Scientific). The DNA was quantified at an average concentration of 150 ng/µL using a Fluoroskan Ascent FL fluorometer (Thermo Fisher Scientific).

### Preparation of the DNA library and sequencing

The DNA library was prepared using DNA extracted from four *A. manihot* accessions. The NEBNext® Ultra™ II DNA Library Prep Kit for Illumina (New England Biolabs, #E7645) was used for DNA library construction according to the manufacturer’s instructions. 500 ng of each DNA extract were sheared separately using mechanical fragmentation on a Bioruptor® (Hologic Diagenode, Belgium) to obtain an average fragment size of 300 bp. The sheared fragments were checked using a 4200 TapeStation with a D5000 screen tape (Agilent Technologies Inc., Santa Clara, CA, USA). After DNA quantification on a Qubit Flex fluorometer (Thermo Fisher Scientific, USA), pooling was performed to obtain at least 200 ng of fragmented DNA input. The quality of the DNA library was checked. Fragments were predominantly between 200 and 800 pb in size, with an average of 335 bp. The DNA library was quantified using the Takara kit (#638,324) on a qPCR machine (LightCycler® 480 Real-Time PCR System, Roche Life Science).

Finally, the library was sequenced using an Illumina MiSeq system (Illumina) provided by the CIRAD genotyping platform, Montpellier, France. A 500 cycle NANO V2 Illumina cartridge (2 x 250 pb) was used to sequence the library in paired-end. A total of 1,295,217 pair-end reads were generated from the DNA library.

### Genome assembly, identification of candidate SSRs and primer design

For the bioinformatic analyses, we used the resources of the SDM-MESO HPC platform at the University of Montpellier (https://isdm.umontpellier.fr).

Sequences were assembled using the ABySS software [9] with a k-mer length of 64 (Table 1). From the obtained assembly, SSRs were detected using the MISA Perl script [10] with following search parameters: at least six repeats for dinucleotide motifs, five repeats for trinucleotide motifs, and five repeats for tetra-, penta- and hexanucleotide motifs. Primers were designed using Primer3 software [11] with default settings. Finally, the primer pairs were blasted with the NCBI Basic Local Alignment Search Tool (BLAST) on the okra (*A. esculentus*) reference genome using the blastn-short parameter of the *blastn* suite [12]. For that purpose, we used the okra genome sequencing data that Wang et al. [13] deposited at the National Genomics Data Center of the China National Center for Bioinformation (https://ngdc.cncb.ac.cn/gwh) under accession number GWHBWBG00000000. The number of hits on the okra genome was obtained from the blast results. We selected the primers pairs showing only one hit. A final selection was made on motif design (from 2 to 6), which resulted in 96 microsatellite loci retained for screening.

### Screening for polymorphic SSR markers

The 96 candidate primer pairs identified were tested for amplification on a subset of 23 *A. manihot* samples. The initial DNA solutions (concentration 150 ng/µL on average) were very viscous, likely due to the presence of sticky, mucilaginous compounds. To facilitate pipetting and reduce the concentration of these compounds that could inhibit the polymerase chain reaction, the DNA solutions were diluted by at least 50-fold. We also had to purify a few recalcitrant samples with AMPure XP magnetic beads (Beckman Coulter, A63881) with a 0.8X bead/DNA ratio. PCR was performed separately for each primer pair in 10 µL final solution, containing 5-10 ng template DNA, 0.5 mM MgCl_2_, 200 µM dNTPs, 0.2 µmol reverse primer, 0.16 µmol M13-tailed forward primer, 0.2 µmol M13 primer fluorescently labelled with FAM, VIC, PET, or NED (Applied Biosystems, Foster City, California, USA), 0.04 ng bovine serum albumin, 0.75X Q-Product (Qiagen, Hilden, Germany) and 0.06 U Taq DNA polymerase with 1X ThermoPol® reaction buffer (New England Biolabs, Ipswich, MA, USA). PCR was performed on a thermocycler Mastercycler® nexus, Eppendorf AG (Hamburg, Germany) with the following settings: initial denaturation at 95 °C for 5 min; 10 cycles of touchdown, starting at 95 °C for 30 s, then annealing temperature (T_a_) plus 5 °C (0.5 °C decrease at each cycle) for 1 min, and 68 °C for 1 min; 23 cycles at 95 °C for 30 s, T_a_ for 1 min, and 68 °C for 1 min; and a final elongation step at 68 °C for 10 min. Allele size was determined after separation on an ABI 3500 xL Genetic Analyzer (Applied Biosystems, Foster City, California, USA). Genemapper v.6.0 software (Applied Biosystems, Thermo Fisher Scientific) was used for data collection, calculation of allele size and visualization of alleles. Primer pairs were classified according to their polymorphism and the overall quality of the profile. Ten of them showed no amplification reaction, 30 loci were monomorphic, 25 loci showed low polymorphism, and 11 loci were difficult to score. Lastly, we selected 21 polymorphic markers (19 mono-locus and one with two different loci) with no ambiguity in allele size determination (Annex 1).

### Validation of the 21 selected microsatellite markers

Genotyping was carried out on 45 accessions of *A. manihot* from three Pacific countries using the 21 SSR markers using the method described above. We used the R package *poppr* [14] to calculate the number of alleles per locus and per country (Table 2), the R package *nikostourvas/PopGenUtils* to calculate the polymorphism information content (PIC) values (Annex 1), and the DARwin software [15] to compute the dissimilarity matrix with simple matching dissimilarity index and perform Principal Coordinate Analysis (Fig. 1).

## CRediT AUTHOR STATEMENT

R. Rivallan: Investigation (sequencing, genotyping), Writing-original draft

A. Garavito: Formal analysis (bioinformatics), Writing-original draft

F. Lawac: Resources, Writing-review & editing

N. Robert: Resources, Writing-review & editing

J. Paofa: Resources, Writing-review & editing

J.-P. Labouisse: Conceptualization, Formal analysis, Writing-original draft, Writing-review & editing, Supervision

## ACKNOWLEDGEMENTS

The authors thank the directions, researchers and technical staff of IAC, NARI, and VARTC who provide the material used in this study. Computations were performed on the ISDM-MESO HPC platform, funded in the framework of State-region planning contracts (Contrat de plan État-région – CPER) by the French Government, the Occitanie/Pyrénées-Méditerranée Region, Montpellier Méditerranée Métropole, and the University of Montpellier. The authors are most grateful to the CIRAD genotyping platform, part of Regional Genotyping Technology Platform, Montpellier, France for its technical support. They thank Dr Vincent Lebot (CIRAD) for funding acquisition and reviewing the article. This work was implemented in the framework of the project No 2271 “*Indications Géographiques pour le Pacifique”* (2022-2024) supported by the *Fonds de coopération économique, sociale et culturelle pour le Pacifique* (in short *Fonds Pacifique*) of the French Ministry of European and Foreign Affairs.

---

**Annex 1.**
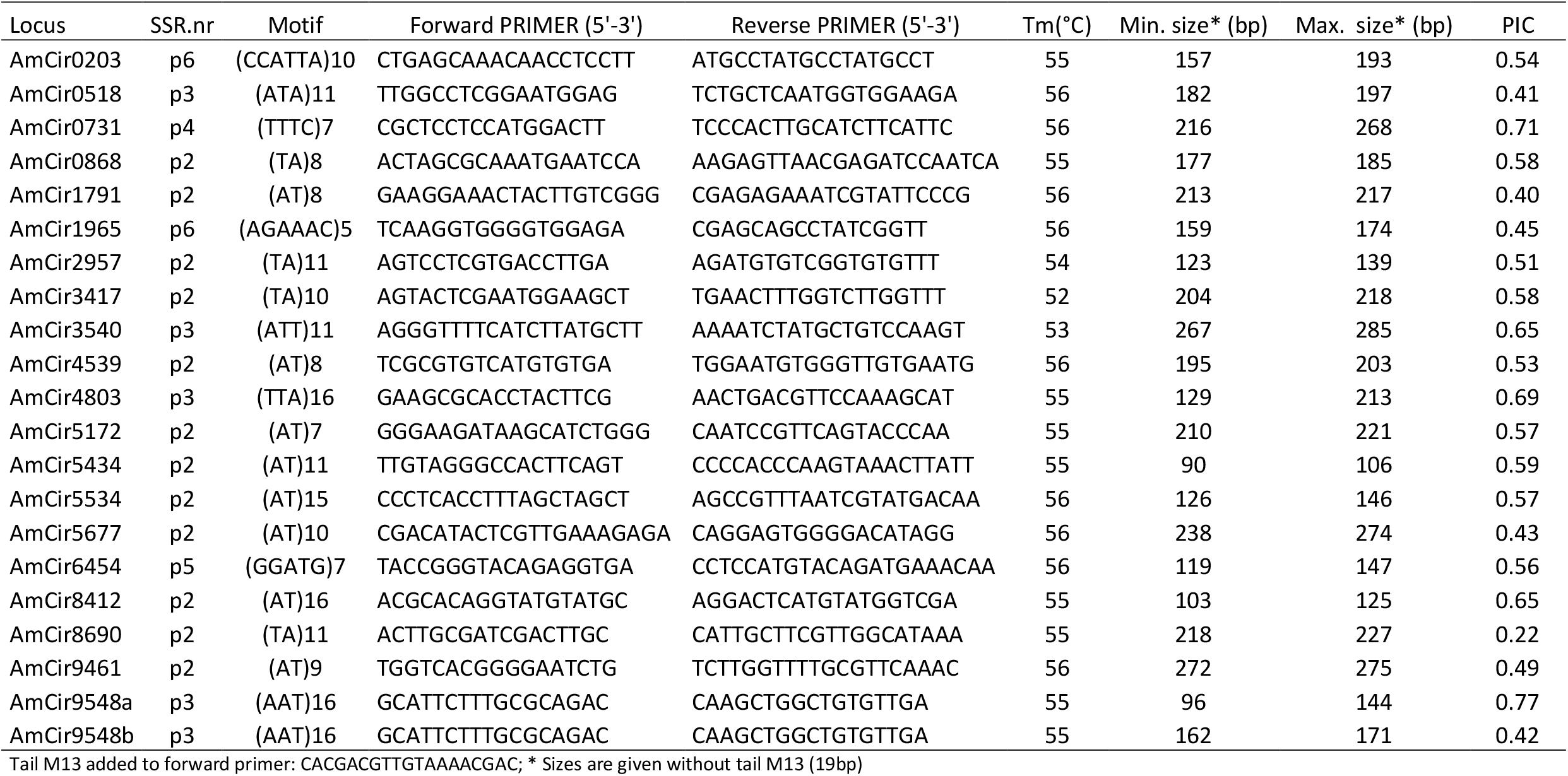
Characteristics of the 21 selected markers of *Abelmoschus manihot*. The minimum and maximum sizes, and the polymorphism information content (PIC) of the markers were obtained from the genotyping results of the 45 accessions.

